# FAMOSS, a conserved 41-aa peptide involved in plant tip growth regulation

**DOI:** 10.1101/2021.11.24.469821

**Authors:** Anna Mamaeva, Andrey Kniazev, Ilia Sedlov, Nina Golub, Daria Kharlampieva, Valentin Manuvera, Victor Rakitin, Alexander Nosov, Artem Fomenkov, Marat Pavlyukov, Sergey Kovalchuk, Rustam Ziganshin, Anna Glushkevich, Vassili Lazarev, Igor Fesenko

## Abstract

Recent evidence shows that small open reading frame (smORF; <100 codons)-encoded peptides (SEPs) containing transmembrane domains are preadapted to be progenitors of novel functional genes. A dozen of such SEPs translated from long non-coding RNAs (lncRNAs) are already functionally characterised in animals. However, functional plant lncRNA-smORF-coded peptides are not yet described. Here, we report detailed functional characterization of a 41-aa peptide encoded by lncRNA-smORFs in the moss Physcomitrium patens, which was named “FAst-growing MOSS” (FAMOSS). We found that the FAMOSS interacts with the Rab-type small GTPase proteins and its overexpression leads to faster moss growth rate and more intensive vesicular transport in apical cells, while its knockout results in the opposite effect. The FAMOSS contains a predicted transmembrane domain and possible orthologs from streptophyta algae to flowering plants have a very conserved structure. Thus, the FAMOSS peptide is a previously unknown conserved player of Rab-mediated processes in plants. Our findings are in line with functional studies of transmembrane SEPs in animals and prove the principles of SEPs evolution. This study provides new insights into functions of plant lncRNA-smORFs.

## Introduction

Eukaryotic transcriptomes contain numerous RNAs with low coding potential (non-coding RNAs), including long non-coding RNAs (lncRNAs), primary microRNAs (pri-miRNA), and circular RNAs (circRNAs) (Li et al., 2017). LncRNAs are transcripts longer than 200 nucleotides that contain only small open reading frames (smORFs; <100 codons) and are unlikely to encode functional proteins. Despite this definition, several functional peptides and microproteins have recently been shown to be encoded by smORFs in mice (Anderson et al., 2015), humans (D’Lima et al., 2017; Huang et al., 2017; Slavoff et al., 2014), zebrafish (Chng et al., 2013), fruit flies (Immarigeon et al., 2021; Magny et al., 2013), *Arabidopsis thaliana* (Hanada et al., 2013), and the moss *Physcomitrium* (previously *Physcomitrella*) *patens* (Fesenko et al., 2019). These smORF-encoded peptides (SEPs) are involved in the regulation of cell proliferation (Polycarpou-Schwarz et al., 2018), cancer growth (Guo et al., 2020; Huang *et al*., 2017), apoptosis and autophagosome formation (Rubtsova et al., 2018), mitochondrial respiratory chain activity (Chugunova et al., 2019), muscle function (Matsumoto et al., 2017), antigen presentation (Niu et al., 2020), heart development (Chng *et al*., 2013), and sexual reproduction (Immarigeon *et al*., 2021). Unlike for animals, only a few examples of functional SEPs have been described in plants. They have been shown to regulate plant programmed cell death (Blanvillain et al., 2011), root growth and development (Casson et al., 2002; Chen et al., 2020), cell proliferation and expansion (Brito et al., 2018; Ikeuchi et al., 2011), nodule formation (Rohrig et al., 2002), and pollen germination (Dong et al., 2013).

Because of their small size, smORFs do not contain functional domains or may include only simple ones, such as a transmembrane α-helix. Indeed, a “transmembrane-first” model of gene birth suggests that peptides with a transmembrane domain may accidently appear in thymine-rich non-genic regions (Vakirlis et al., 2020). A pool of uncharacterized SEPs located at, and interacting with, cellular membranes was detected in different studies; for example, the 56-aa Mtln peptide interacts with NADH-dependent cytochrome b5 reductase (Chugunova *et al*., 2019), the 90-aa SPAR peptide interacts with V-ATPase (Matsumoto *et al*., 2017), and the 28-aa Sarcolamban A interacts with Sarco-endoplasmic Reticulum Ca^2+^ adenosine triphosphatase (SERCA) (Magny *et al*., 2013). The identification of protein partners of the SEPs helps elucidate their functions (Matsumoto *et al*., 2017; Slavoff *et al*., 2014); for example, a human 7-kDa microprotein, NoBody, binds the mRNA decapping complex to regulate the number of P-bodies in a cell (D’Lima *et al*., 2017). The 10-kDa microprotein CASIMO1 controls cell proliferation by interacting with squalene epoxidase that is a key enzyme in cholesterol synthesis and a known oncogene in breast cancer (Polycarpou-Schwarz *et al*., 2018).

Previously, we identified functional smORFs encoded by lncRNAs in the model plant *Physcomitrium patens*, a moss (Fesenko *et al*., 2019). The knockout of four selected SEPs, PSEP1, PSEP3, PSEP18, and PSEP25, resulted in significantly reduced growth rates in the moss protonemata (Fesenko *et al*., 2019). In this study, we investigated the functions of PSEP1, which was renamed the FAst-growing MOSS (FAMOSS) peptide. We demonstrated that this SEP participates in the regulation of polar tip growth in *P. patens* via its interaction with Rab-type small GTPases. We observed that mutant plants lacking FAMOSS had altered apex-derived vesicular trafficking and different protonemal filament structures compared with the wild type. Our quantitative proteomic analysis revealed a downregulation of the stress-responsive proteins in both the *FAMOSS-*overexpressing (OE) and knockout (KO) lines, as well as an altered sensitivity to phytohormones. Finally, we proposed that the FAMOSS peptide is a novel component of the Rab protein complexes that regulate vesicular trafficking and stress responses in plants.

## Results

### FAMOSS is widely distributed across different plant lineages

Recently, we used a combined approach to identify functional SEPs encoded by lncRNAs in the model plant *P. patens* (Fesenko *et al*., 2019). In this analysis, we identified the 41-aa peptide PSEP1 (Physcomitrella smORF-encoded peptide 1), encoded by the transcript CNT2064811, which was predicted to be a non-coding RNA in the CANTATAdb database (Szczesniak et al., 2016) but was missed in the Phytozome V12 annotation (Goodstein et al., 2012; Lang et al., 2018). The overexpression of this peptide resulted in larger moss plant sizes and a faster growth rate, whereas the KO mutant had the opposite phenotype (Fesenko *et al*., 2019). Based on the observed phenotypic effects, we renamed this peptide “FAMOSS” here.

The inspection of the genome region around CNT2064811 revealed that the protein-coding gene *Pp3c9_20610V3* is located in close proximity to this lncRNA. To exclude the possibility that the FAMOSS peptide is an upstream or downstream smORF located within annotated mRNA, we additionally performed direct nanopore RNA sequencing (ONT DRS) on the protonemal and gametophore transcriptomes (Fesenko et al., 2021). ONT DRS allows the accurate assembly and identification of the exon–intron structure of the transcripts. This analysis confirmed that CNT2064811 is an independent transcriptional unit with a higher transcriptional level than the surrounding protein-coding genes (Figure 1A). To additionally explore whether FAMOSS is translated from the predicted smORF, we used CRISPR-Cas9 to fuse a reporter *GUS* sequence to the *FAMOSS* sequence in the moss genome (Figure 1B–D). We observed that *FAMOSS* expression both in the protonemata and gametophores was consistent with the results of the transcriptomic analysis.

**Figure 1.**
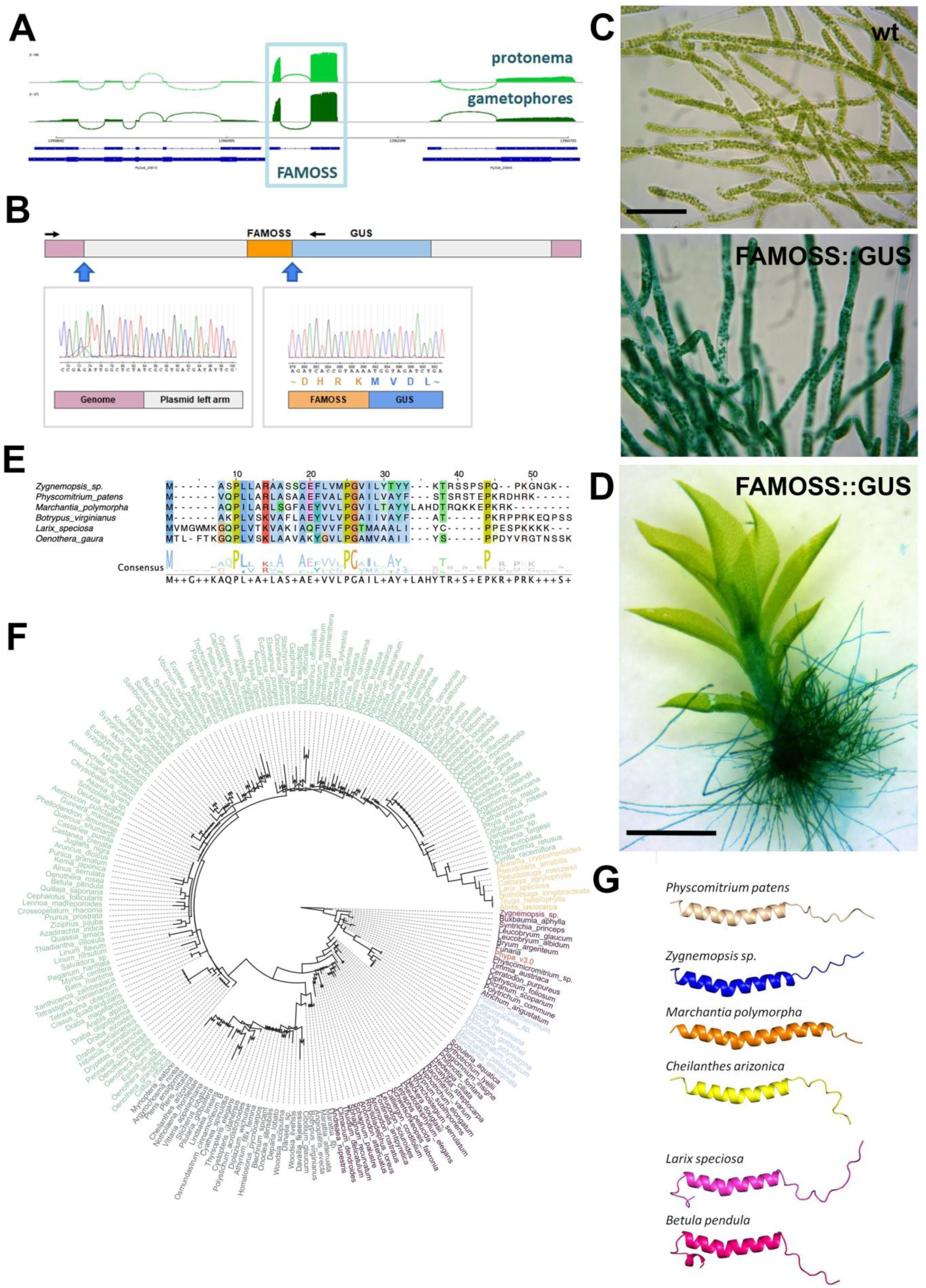
FAMOSS is a widely conserved plant smORF-encoded peptide. A - Sashimi plot showing *FAMOSS* transcription as confirmed using nanopore RNA sequencing in *Physcomitrium patens* protonema and gametophores. B - Construction of FAMOSS coupled with a GUS reporter protein at the FAMOSS promoter. Black arrows indicate primers used for identification. C, D - GUS staining of the protonema (C, scale bar = 200 μm) and gametophores (D, scale bar = 1 mm). E - The alignment of the FAMOSS peptide and orthologous sequences from different plants. F - Phylogenomic tree of the identified FAMOSS orthologs. Mosses, burgundy; liverworts, blue; algae, red; ferns, gray; gymnosperms, orange; angiosperms, green. G - *P. patens* FAMOSS structure and structures of its orthologs, predicted using AlphaFold.

We next analyzed the conservation of the FAMOSS peptide across different plant lineages. Using a peptide amino acid sequence as a query, we performed a TBLASTN search (*E*-value cutoff < 0.001) against the transcriptomes from the 1000 plants project (OneKP) and manually inspected the results. This search revealed possible orthologs in the liverworts, other mosses, and ferns, suggesting that this peptide was widely conserved in certain lineages (Figure 1E, F); however, no hits could be detected in the hornworts, lycophytes, or vascular plants. In addition, an ortholog of the FAMOSS peptide was also found in the transcriptome of *Zygnemopsis* sp. (streptophyte algae). Although the C-terminal end of the FAMOSS peptide was poorly conserved, the analysis of multiple alignments between the possible orthologs in green algae, mosses, liverworts, and ferns revealed a pattern of highly conserved amino acid residues (Figure 1E). To identify potential conserved motifs in the FAMOSS peptide, the MEME algorithm (Bailey et al., 2015) was used. This approach confirmed the presence of a significantly conserved motif, spanning the first 30 aa of the FAMOSS peptide (Supplemental Figure 1).

Given that the identification of peptide homologs can be problematic in distant species, we next performed a HMMER search (*E*-value cutoff < 0.01) against the translated transcriptomes of selected Gymnospermae and Angiosperms species from the OneKP project (One Thousand Plant Transcriptomes, 2019) and manually curated the results. This analysis revealed possible orthologs in eight Gymnospermae species (12 transcripts, mean = 44 aa), 108 Core Eudicots/Rosids species (124 transcripts, mean = 47 aa), 41 Core Eudicots/Asterids species (53 transcripts, mean = 53 aa), and 25 Basal Eudicots (38 transcripts, mean = 46 aa) (Supplemental Table 1); however, further studies are needed to confirm that these peptides are true orthologs. The corresponding transcripts do not encode ORFs longer than 100 aa, and 50% (63/124) of the transcripts from the Core Eudicots/Rosids, 66% (35/53) from the Core Eudicots/Asterids, and 71% (27/38) from the Basal Eudicots were annotated as lncRNAs in a database of lncRNAs in Angiosperms, AlnC (Singh et al., 2021).

According to a prediction made using TMHMM version 2.0 (Krogh et al., 2001), the *P. patens* FAMOSS peptide contains a transmembrane domain (Fesenko et al., 2021). To further investigate the relationship between the conserved amino acid residues and the structural properties of FAMOSS, we predicted the *P. patens* FAMOSS 3D structure using a recently developed machine learning algorithm, AlphaFold2 (Jumper et al., 2021), which was shown to have promising results. According to the obtained structural model, the N-terminal and central parts of the peptide represent an alpha-helix with a slight kink in its center, probably associated with a proline residue. The C terminus was predicted to be disordered (Figure 1G). To further explore this finding, we analyzed the structures of predicted FAMOSS orthologs from five other plants from different taxonomic groups (algae, liverworts, ferns, conifers, and dicots), revealing similar results overall (Supplemental Figure 2). Our findings suggest that an ortholog of the FAMOSS peptide was present in the last common ancestor of land plants and streptophyte algae and widely conserved in different land plant lineages. The high expression level of the corresponding transcript suggests that this peptide is an unannotated highly conserved small protein.

### FAMOSS regulates the tip growth rate in protonemata

Previously, we showed that *P. patens FAMOSS* OE mutant lines displayed a rapid protonemal growth phenotype, whereas the KO lines grew more slowly than the wild-type plants (Fesenko *et al*., 2019). Moss protonemata consist of two morphologically distinguishable cell types, chloronemata and caulonemata (Figure 2A). Chloronemata are produced first, as the spores germinate, while caulonemal cells develop several days later at the tips of the protonemata (Vidali and Bezanilla, 2012). Caulonemal cells are characterized by their higher growth rate than chloronemal cells and tangential cell walls (Jang and Dolan, 2011).

**Figure 2.**
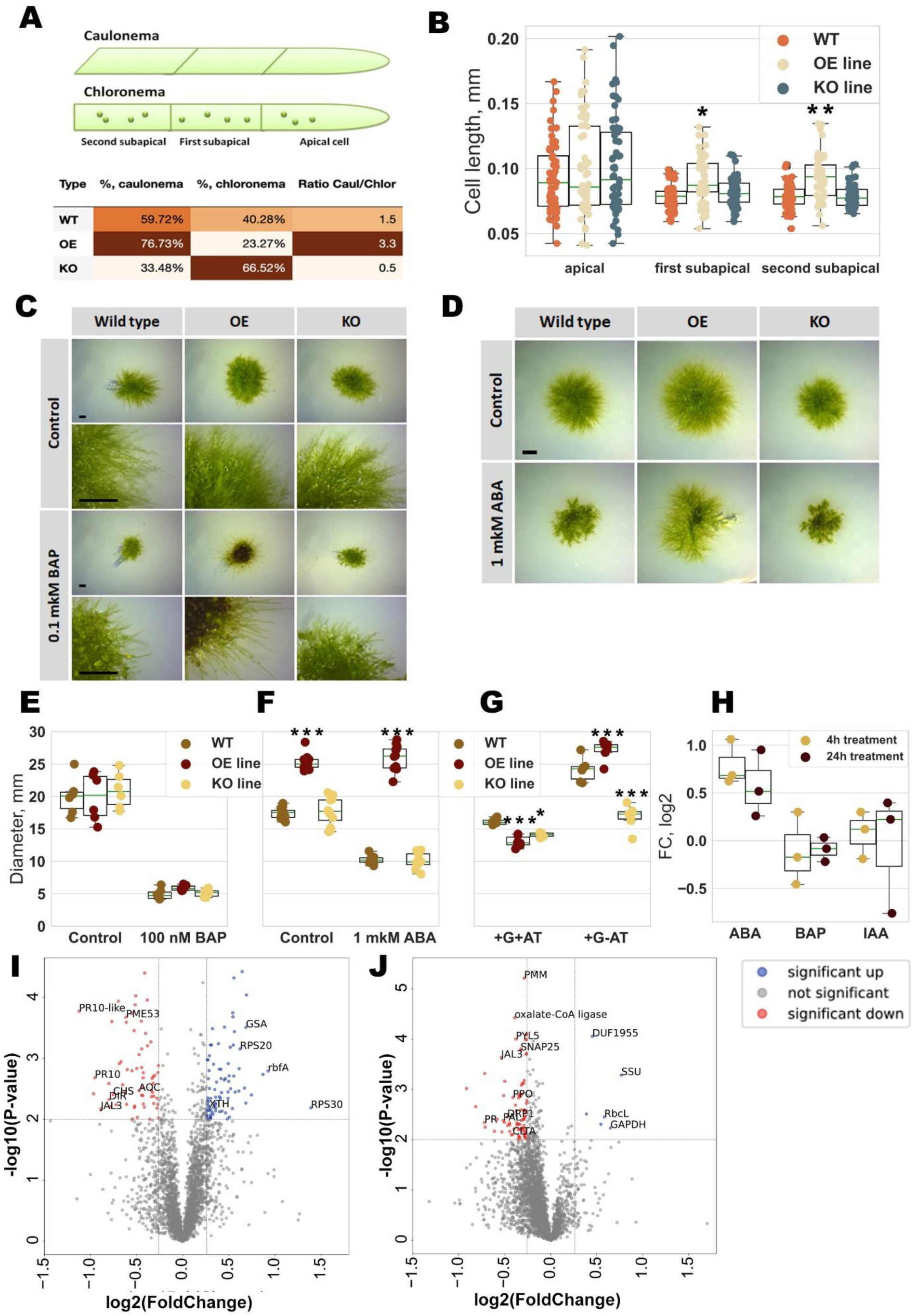
The growth of mutant lines in different conditions. A - The ratios of the two types of moss protonema filaments, caulonemata and chloronemata, in wild-type, *FAMOSS-*overexpressing (OE), and knockout (KO) plants. B - The lengths of the apical, first subapical, and second subapical cells. C - The effect of 6-benzylaminopurine (BAP) on protonemal morphology. Scale bar = 1 mm. D – The effect of abscisic acid (ABA) on protonemal morphology. E - The effect of BAP on the protonemal growth rate. F - The effect of ABA on protonemal growth. G - The effect of medium composition on protonemal growth. +G+AT, BCD medium supplemented with glucose and ammonium tartrate; +G–AT, BCD medium with glucose but without ammonium tartrate. H – Fold change (FC) in *FAMOSS* transcription after treatments with ABA, BAP, and indole-3-acetic acid (IAA). I, J - Quantitative proteomic analysis of the *FAMOSS*-overexpressing (OE) and knockout (KO) plants. Volcano plots of the entire set of proteins quantified in the iTRAQ analysis in the *FAMOSS* OE (I) and KO (J) lines. Proteins with significant differences in abundance relative to the wild type are depicted in color. GSA, glutamate-1-semialdehyde 2,1-aminomutase; rbfA, ribosome-binding factor A; RPS20, ribosomal protein S20; RPS30, ribosomal protein S30; AOC, allene oxide cyclase; GST, glutathione *S*-transferase; CHS, chalcone synthase; XTH, xyloglucan endotransglucosylase/hydrolase protein 23; PME, pectin methylesterase; rbcL, ribulose-1,5-bisphosphate carboxylase/oxygenase large subunit; GAPDH, glyceraldehyde-3-phosphate dehydrogenase A chloroplastic-like isoform X2; JAL3, jacalin- related lectin 3; PPO, polyphenol oxidase; PAL, phenylalanine ammonia-lyase; PR10, pathogenesis-related protein; PR10-like, pathogenesis-related protein-like; DIR, dirigent protein; SNAP25, synaptosomal-associated protein 25; ASPL, tether containing UBX domain for GLUT4; DRP1, dynamin-related protein 1E; CLTA, clathrin light chain; PYL5, pyrobactin resistance 1-like 5; PMM, phosphomannomutase; SSU, ribulose-1,5-bisphosphate carboxylase/oxygenase small subunits; DUF1955, domain of unknown function. *, *P* < 0.05; **, *P* < 0.01; ***, *P* < 0.001.

To test the hypothesis that the rapid growth of the *FAMOSS* OE lines is due to the accumulation of caulonemal filaments, the proportion of chloronemal to caulonemal cells was compared in the wild-type and mutant lines. The ratio of caulonemal to chloronemal cells was significantly higher in the OE line and lower in the KO line when compared with the wild type (Figure 2A; *n* = 3 independent biological repeats; chi-square test, *P* < 10^−15^). The chloronemal cells therefore predominate in the KO line (Supplemental Figure 3), suggesting a link between *FAMOSS* expression and the transition from chloronemal to caulonemal cells in *P. patens*. We next compared the length of the apical and two subapical cells between the wild-type and mutant lines. Although the lengths of the apical cells were similar in all genotypes, the subapical cells were significantly longer in the OE line than in the wild type or the KO line (Figure 2B, ANOVA with post-hoc Tukey HSD (Honestly Significant Difference) *P* < 0.00001). These observations led to the hypothesis that the observed changes in the moss growth rate come from an altered protonemal structure (caulonemata/chloronemata ratio) and increased subapical cell lengths.

The chloronemal-to-caulonemal transition is regulated by both hormonal crosstalk and different factors, such as glucose, light, and low nutrient availability (Jaeger and Moody, 2021). Low light, cytokinins, and abscisic acid (ABA) are negative regulators of caulonemal differentiation, whereas auxin, glucose, and a high light intensity are positive regulators. We therefore investigated how both negative and positive regulators influence the caulonemal transition in the wild-type and mutant lines (Fig. 2C–F).

First, we tested the influence of cytokinin and ABA treatments on the growth rate of both mutant lines (Figure 2C, D). It has previously been shown that large concentrations of cytokinins in the growth medium inhibit filament growth and induce callus-like structures (Sabovljevic et al., 2014). In our experiments, filament growth in the wild-type and KO plants was strongly inhibited under a 0.1 μM 6-benzylaminopurine (BAP) treatment, but the *FAMOSS* OE mutant line produced long protonemal filaments, suggesting a decreased sensitivity to cytokinins (Figure 2C, E). Another negative regulator, ABA, significantly inhibited the growth of both the wild-type and KO moss (Figure 2D and F; ANOVA with post-hoc Tukey HSD *P* < 0.001), but there was no effect on the growth rate of the OE line. Thus, the overexpression of *FAMOSS* led to a reduced sensitivity to the negative regulators of moss protonemal growth, cytokinin, and ABA.

Ammonium tartrate (AT) is known to reduce the transition into caulonemal cells and promote chloronemal branching (Vidali and Bezanilla, 2012). Given the altered ratios of chloronemal to caulonemal filaments in the mutant lines, we compared the growth of the different genotypes on culture mediums with and without AT. The application of AT significantly reduced the growth rates of both the wild-type and mutant plants (*P* < 0.01; Figure 2G), although both mutant lines significantly differed from the wild type, with the OE plants being the smallest. On the medium without AT, the OE plants showed the fastest growth, but the KO plants grew more slowly than the other genotypes (Figure 2G).

The stimulatory effect of auxin on the transition from chloronemal to caulonemal filaments has been shown previously; however, exogenous auxin treatments resulted in the inhibition of moss growth (Thelander et al., 2018). In our experiments, the growth of wild-type moss plants was significantly inhibited under auxin treatment starting with 1 nM indole-3-acetic acid (IAA; Supplemental Figure 4). In contrast, the growth rate of the OE line was significantly decreased only by 1 μM IAA (ANOVA with post-hoc Tukey HSD *P* < 0.001). As the auxin concentration increased, the protonemal filaments were increasingly composed of caulonemata, resulting in a sparser and lighter appearance. Thus, *FAMOSS* is expressed in conditions favorable to caulonemata formation, causing chloronemata to differentiate into caulonemata, which increases the protonemal growth rate.

We next tested whether the auxin and cytokinin treatments influenced the transcriptional level of *FAMOSS*; however, no difference was observed after a 4- or 24-h treatment with either phytohormone (Figure 2H). By contrast, *FAMOSS* expression was marginally upregulated by ABA (two-tailed t-test, *P* < 0.05). Our findings showed that the overexpression of *FAMOSS* positively regulates protonemal growth and the transition from chloronemal to caulonemal filaments. In particular, sensitivity to the hormonal negative regulators of the caulonemal transition was reduced in the OE line. In addition, the proportion of caulonemal filaments was reduced in the KO plants, which had a similar phenotype to the wild-type plants, although the growth rate was lower on the medium without the inhibitor of caulonemal growth (AT).

### Proteomic analysis of the mutant lines

We next used isobaric tags for relative and absolute quantitation (iTRAQ)-based comparative quantitative proteomic analyses to identify changes in the proteomes of the mutant lines. Because iTRAQ quantification underestimates the amount of real fold change (Ow et al., 2009), fold change ratios of >1.20 or <0.83 (*P* < 0.01, one-way ANOVA) were used to identify differentially expressed protein groups (DEPs). Overall, we identified 118 DEPs in the OE line and 75 DEPs in the KO line (Figure 2I, J; Supplemental Table 2).

The most upregulated DEPs in the OE line were ribosomal proteins, such as Pp3c2_29010 (RP-S30e), Pp3c24_17080 (RP-S20), and Pp3c12_16410 (rbfA) (Figure 2I). Among the most downregulated DEPs in the OE line, we identified a group of proteins that participate in the stress response; for example, the pathogenesis-related proteins (PR-10) Pp3c2_27350 and Pp3c7_19850 were previously shown to be involved in the defense response (Castro et al., 2016) and downregulated under osmotic stress (Stevenson et al., 2016). In addition, proteins related to the biotic stress response were downregulated, including Pp3c5_22560 (dirigent protein; DIR), Pp3c5_22560 (chalcone synthase; CHS), and Pp3c4_22490 (allene oxide cyclase; AOC). Stress-induced DIR plays a role in the adaptive response (Paniagua et al., 2017), while CHS is a key enzyme of the flavonoid/isoflavonoid biosynthesis pathway and is involved in the salicylic acid biosynthesis pathway (Dao et al., 2011). AOC is involved in the jasmonic acid biosynthesis pathway (Stenzel et al., 2003). Another protein related to secondary metabolism regulation and the defense response is Pp3c9_1620 (polyphenol oxidase; PPO) (Tran et al., 2012). The protein pectin methylesterase (Pp3c5_12660) was also downregulated in the OE line. Pectin methylesterases modulate the cell wall mechanical properties and organ initiation in Arabidopsis (Peaucelle et al., 2011). These findings therefore suggest that the OE line would be less resistant to a variety of stress conditions. However, additional experiments are needed to confirm or dismiss this hypothesis.

The proteome changes in the KO line were less pronounced (Figure 2J). The chloroplastic proteins, such as the large and small subunits of RuBisCO and Pp3c11_15790 (glyceraldehyde 3-phosphate dehydrogenase (GAPDH)), were among the most upregulated in the KO line (Supplemental Table 2). One possible explanation for this result is the predominance of chloroplast-rich chloronemal filaments in this line. The stress-related proteins were also downregulated in the KO line, as was seen in the proteome of the OE line; for example, Pp3c7_19850 (PR10), Pp3c6_6545 (DIR-related), and Pp3c1_18940 (phenylalanine ammonia-lyase; PAL) were downregulated. It is worth noting that Pp3c7_19850 (PR10), Pp3c18_10760 (glutamine synthetase), Pp3c9_1620 (polyphenol oxidase), and Pp3c1_1940 (jacalin-like lectin domain) were downregulated in both the OE and the KO lines. The top 10 most downregulated proteins included two proteins containing the cupin (Cupin_2) domain (Supplemental Table 2). Proteins with this domain play crucial roles in plant development and defense (Wang et al., 2014).

We also identified a group of downregulated DEPs associated with vesicle trafficking in the KO line. These were dynamin-related protein 1C (DRP1C, Pp3c19_4870), which regulates vesicle formation in post-Golgi trafficking (Fujimoto and Tsutsumi, 2014; Jilly et al., 2018); clathrin light chain (Pp3c7_10220; CTLA); and synaptosomal-associated protein-related (Pp3c2_7850; SNAP25). SNAP25 is a SNARE (soluble *N*-ethylmaleimide-sensitive factor attachment protein receptor) protein, which is involved not only in vesicular trafficking under normal conditions but also in regulating transport of cell wall–associated and defense proteins during biotic stress (Kwon et al., 2020). SNARE-mediated membrane trafficking is also required for the proper response of plants to abiotic stresses.

The iTRAQ analysis thus showed that proteins involved in the response to different stress conditions were downregulated in both mutant lines. By contrast, the most upregulated DEPs in the OE line belong to biosynthetic processes, suggesting that the rapid growth rate in the OE line is linked to proteome changes.

### The stress response is altered in the mutant lines

According to the quantitative proteomic analysis, some stress-related proteins were downregulated in both mutant lines, suggesting that they may have altered stress tolerances. In addition, the reduced sensitivity to ABA observed in the OE line could decrease its resistance to abiotic stress factors.

We next investigated the growth of both the mutant and wild-type plants under abiotic stress conditions induced by salt (NaCl) and paraquat (PQ) (Figure 3A-C). All genotypes were severely affected starting with the 150 mM NaCl concentration (Figure 3A); however, the growth rate of the OE plants was significantly reduced compared with the wild type and KO under this concentration of NaCl (*P* < 0.001, ANOVA with post-hoc Tukey HSD test). Moreover, the OE line had the most severely altered phenotype after the 150 mM NaCl treatment (Figure 3C).

**Figure 3.**
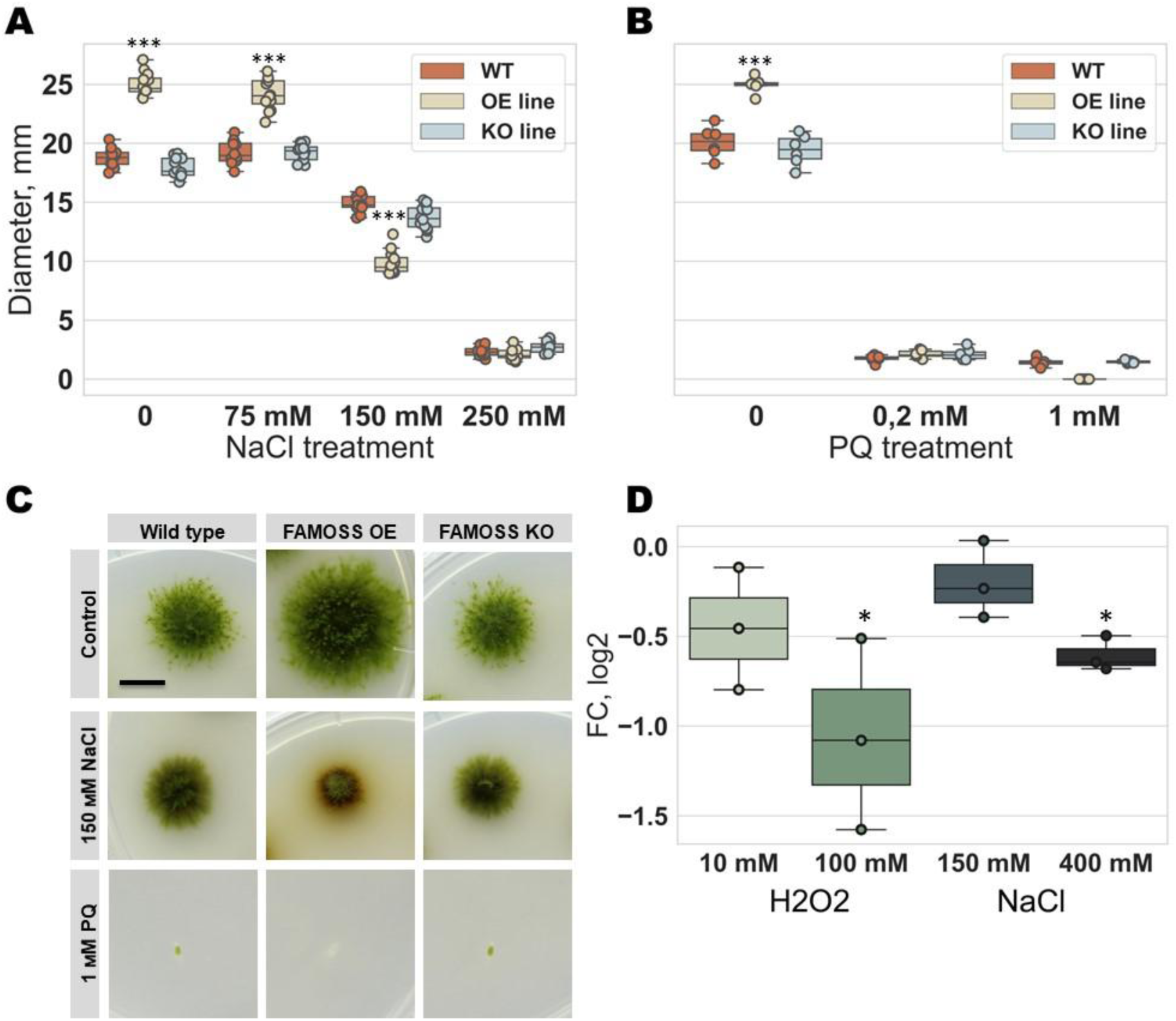
The growth rate of the mutant lines under stress-related conditions A - Effect of NaCl on protonemal growth. B - Effect of paraquat (PQ) on protonemal growth. C - Effect of NaCl and PQ on protonemal morphology after 30 days of cultivation. D - *FAMOSS* expression under different stresses. *, *P* < 0.05; ***, *P* < 0.001.

PQ is a potent reactive oxygen species (ROS) inducer. The growth of all genotypes was significantly inhibited by a treatment with either 0.2 or 1 mM PQ (*P* < 0.001, ANOVA with post-hoc Tukey HSD test; Figure 3B, C). Moreover, the growth of the OE line was completely inhibited and the plants died under 1 mM PQ (Figure 3C).

Using RT-qPCR analysis, we found that the level of *FAMOSS* transcription in the wild-type protonemata was only marginally decreased after a 1-h treatment with 400 mM NaCl or 100 mM H_2_O_2_ (two-tailed t-test, *P* < 0.05; Figure 3D). This result suggests that *FAMOSS* is not stress-inducible at the transcriptional level.

Our results showed that the mutants, especially the OE line, had a reduced ability to adapt to stress. Given that ABA is the most important regulator of the abiotic stress response, the rapid protonemal growth and altered ABA sensitivity in the *FAMOSS* OE line could result in the observed phenotypes.

### FAMOSS interactome

We further investigated the possible protein interaction partners of the FAMOSS peptide using two approaches, a pull-down assay and coimmunoprecipitation (co-IP) (Figure 4A-C). For the pull-down assay, a recombinant FAMOSS peptide fused with streptavidin at the N-terminal end (FAMOSS-SAV; see Materials and Methods) and recombinant streptavidin were used (Figure 4B). We used reversible cross-linking to bind the interacting proteins and identified them using a mass spectrometry (MS) analysis. This approach revealed 323 unique protein groups in the FAMOSS-SAV samples (Supplemental Table 3). Besides the large clusters of ribosomal and chloroplast proteins, which are common contaminants (Mellacheruvu et al., 2013), the most promising protein partners were Rab-type small GTPases (Figure 4D). Rab proteins participate in a diverse range of cellular processes, including vesicular trafficking, tip growth, and the stress response (Chen, Heo, 2018; Lycett, 2008; Ma, 2007). Because many contaminants were observed in the FAMOSS-SAV interactomes, we performed a gel electrophoresis analysis of the samples and analyzed the 25-kDa band using an in-gel trypsin digestion followed by a liquid chromatography (LC)–MS/MS analysis to enrich the fraction of Rab proteins.

**Figure 4.**
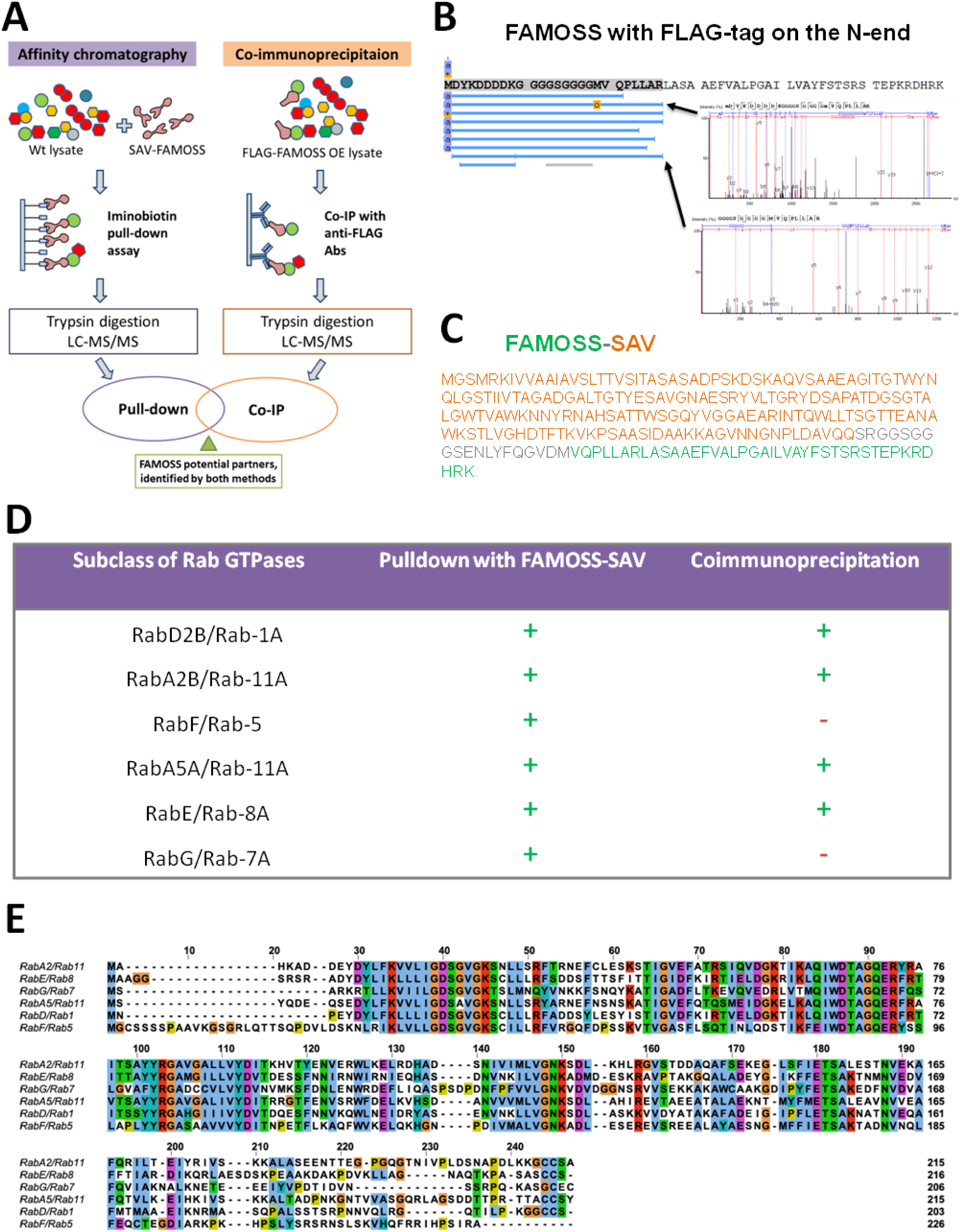
Analysis of the protein partners of the FAMOSS peptide. A - Diagram of the interactome experiments. B - Sequence and coverage of the FLAG-tagged FAMOSS sequence. C - Sequence of streptavidin (SAV)-tagged FAMOSS. D - Rab GTPases present in the two types of interactome experiments. Proteins detected in samples are marked with a plus sign, and those not detected are indicated with a minus sign. E – Alignment of Rab GTPases found in the FAMOSS interactomes.

The final list of Rab-type GTPases, unique for FAMOSS-SAV samples, included RabD2B/Rab-1A, RabA2B/Rab-11A, RabF/Rab-5, RabA5A/Rab-11A, RabE/Rab-8A, and RabG/Rab-7A (Figure 4D; Supplemental Table 3). Among these proteins, the largest peptides coverage (33%) and intensity were observed for the Pp3c14_15250 (RabA2B/Rab-11A) protein.

To validate the obtained results, we generated mutant lines overexpressing FAMOSS*-*FLAG fusion constructs (Figure 4A). The FLAG tag was placed on the C terminus (mutant 1028 line) or on the N terminus (mutant 1032 line) of the FAMOSS peptide (Figure 4B). We immunoprecipitated FLAG-tagged FAMOSS and identified RabD2B/Rab-1A, RabA2B/Rab-11A, RabA5A/Rab-11A, and RabE/Rab-8A in the interactome of the 1032 line (*n* = 3 independent experiments) (Figure 4D). These proteins were unique for the FAMOSS-FLAG interactomes and were not found in the control samples.

Due to high sequence similarity of Rab GTPases (Figure 4E; Lycett, 2008; Ma, 2007; Minamino and Ueda, 2019), we cannot rule out the possibility that FAMOSS peptide interacts with several Rab proteins. For example, human GDI1 protein was shown to interact with different Rab GTPases: RAB3A, RAB5A, RAB5B, RAB5C, RAB8A, RAB8B, RAB10, RAB35 and RAB43 (Steger et al., 2017). In addition, due to similar subcellular localization (Elliott et al., 2020) Rab proteins from different families could co-purify together in our experiments.

Thus, our results suggest that the Rab GTPase proteins may interact with the FAMOSS peptide. The Rab GTPase proteins target vesicles to the plasma membranes and mediate their fusion, thereby stimulating the intensity of vesicular trafficking to the cell apex and increasing the tip growth rate (Lycett, 2008; Ma, 2007). This is consistent with the phenotype of the *FAMOSS* OE lines.

### FAMOSS regulates vesicular trafficking

The moss protonemal filaments expand by polarized tip cell growth similar to pollen tubes or root hairs in flowering plants (Orr et al., 2020; Rensing et al., 2020; Rounds and Bezanilla, 2013; Vidali and Bezanilla, 2012). It is well known that plant tip growth requires active apical vesicular trafficking regulated by different families of small GTPases, including Rab proteins (Minamino and Ueda, 2019). The RabA GTPases were previously shown to regulate polarized secretion and vesicle delivery in plants (Chen, Heo, 2018; de Graaf et al., 2005; MartiniEre and Moreau, 2020; Szumlanski and Nielsen, 2009).

To determine whether the FAMOSS peptide influences the vesicle pool in the apex cells, we labelled protonemal filaments of the wild type and both mutant lines using the fluorescent dye SynaptoGreen C4 (also known as FM1-43, a trademark of Molecular Probes) (Jelinkova et al., 2010) (Figure 5A–C). The FM1-43 dye is widely used for a marking of vesicles in the tip growing plant cells such as pollen tube, root hair and protonema apical cells (Samaj et al., 2005; Bove et al., 2008; Pleskot et al., 2012; Gisbergen et al., 2020). In the wild-type cells, the distribution of the apical vesicles was similar to that which was previously described in *P. patens* (Rawat et al., 2017; Gisbergen et al., 2020) (Figure 5B); however, the labelling intensity was significantly increased in the OE line compared with both the wild-type and KO lines (ANOVA, *P* < 0.001; Figure 5D, E). This suggests a more intensive vesicle trafficking in the cell apex, which is consistent with the rapid protonemal growth in the OE line. By contrast, the fluorescence intensity was significantly decreased in the cell apexes of the protonemal filaments in the KO line compared with the wild-type and OE line cells (ANOVA, *P* < 0.001; Figure 5E).

**Figure 5.**
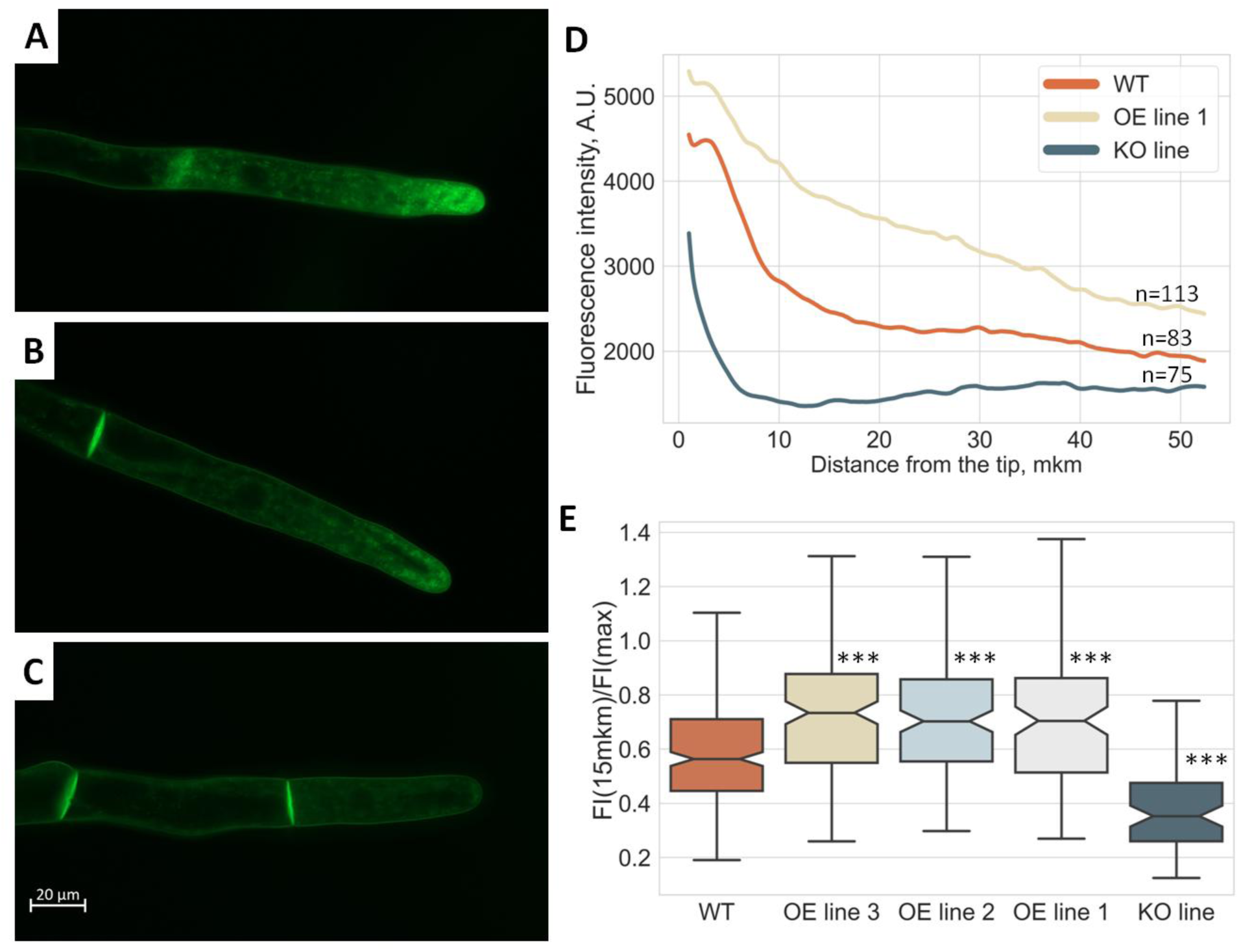
The FAMOSS peptide affects vesicular trafficking. A–C - Apical protonemal cells stained with SynaptoGreen C4. (A – *FAMOSS-*overexpression line (OE); B – The wild type; C - *FAMOSS* knockout line (KO). D - Profile of SynaptoGreen C4 fluorescence in the apical cells. E - Boxplot showing the ratios of intensity of SynaptoGreen C4 staining at the distance 15 mkm from the cell tip to the maximal intensity. ***, *P* < 0.001.

**Figure 6.** The mechanism of the regulation of plant physiological processes by the FAMOSS peptide. *FAMOSS* is expressed in favorable conditions and modulates the Rab signaling pathway to influence vesicular trafficking and ABA signaling. These processes increase protonemal growth, translation, and caulonemal formation, but decrease the ABA response and, thus, the adaptability to stresses.

These results suggest that the FAMOSS peptide may function as a possible modulator of vesicular trafficking during tip cell growth via its interaction with the Rab small GTPases.

Given that apical vesicular transport is linked to cell wall formation (Rounds and Bezanilla, 2013), we next analyzed the regeneration of the protonemal filaments after protoplast isolation. The proportion of regenerated plants with new chloronemal filaments was significantly higher in the KO line (∼18%) than in the wild type (∼8%) or the OE line (∼2%; chi-squared test *P* < 10^−10^). The FAMOSS peptide therefore appears to inhibit the process of chloronemal formation, which results in a more effective regeneration process in the KO line. This correlates with the observed chloronemata/caulonemata ratio in the protonemal tissue.

## Discussion

A growing body of evidence suggests that the peptides or microproteins encoded by smORFs may play important roles in animal and plant cells (Casson *et al*., 2002; Huang *et al*., 2017; Matsumoto *et al*., 2017; Polycarpou-Schwarz *et al*., 2018; Rohrig *et al*., 2002; Slavoff *et al*., 2014; Tavormina et al., 2015); however, there are few examples of functional SEPs in plants. In our previous study, we identified a new 41-aa sORF-encoded peptide that influences moss growth located on a transcript predicted to be a lncRNA (Fesenko *et al*., 2019).

In this study, we identified possible FAMOSS orthologs in liverworts, mosses, ferns, conifers, and eudicots, indicating that it is widely conserved across the plant kingdom. A FAMOSS ortholog was also found in *Zygnemopsis* sp. (streptophyte algae). This is consistent with the findings that the Zygnematophyceae appear to be the closest relatives of land plants (Delwiche and Cooper, 2015; Nishiyama et al., 2018; One Thousand Plant Transcriptomes, 2019). The corresponding transcripts do not encode longer (>100 aa) proteins, suggesting that the *FAMOSS* orthologs can be considered the main ORFs. The conservation of the *FAMOSS* peptide across different plant lineages is consistent with our results, in which a set of Rab-type small GTPases were identified in the FAMOSS interactomes (Table 1). Rab GTPases are highly conserved regulators of membrane trafficking in plants (Lycett, 2008; Ma, 2007; Minamino and Ueda, 2019) and are present in both algae (Hoepflinger et al., 2013) and land plants (Minamino and Ueda, 2019). In particular, proteins from the RabA clade are involved in the coordination of tip cell growth (Ma, 2007); for example, these proteins are crucial for the tip-targeted vesicular trafficking of cell wall components in root hairs (Preuss et al., 2004) and pollen tubes (Szumlanski and Nielsen, 2009) in Arabidopsis. Tip growth of the moss protonemata is also regulated by the Rab proteins (Orr et al., 2021); thus, the consideration of Rab GTPases as possible partners of the FAMOSS peptide is consistent with the protonemal growth rates, subapical cell lengths, and chloronemata-to-caulonemata transition in the mutant lines. Moreover, using the fluorescent dye SynaptoGreen C4, we revealed changes in the intensity of the vesicular trafficking in the mutant lines. A previous study showed that the AtRabA1d protein accumulated in the apical dome of the growing root hairs in Arabidopsis, completely colocalizing with intracellular vesicles stained with the fluorescent dye FM4-64 (Ovecka et al., 2010). In addition, the quantitative proteomic analysis revealed that some proteins associated with vesicular trafficking were downregulated in the KO plants. Considering results from different experiments, we hypothesize that the interaction of the FAMOSS peptide with Rab GTPases or with proteins from the corresponding protein complex may affect the intensity of vesicular trafficking in the moss tip cells and influence the protonemal growth rate.

In addition to regulation of tip growth in plants, Rab proteins are also involved in phytohormone signaling and the stress response. AtRabE1c interacts with the ABA receptor PYL4 (pyrobactin resistance 1-like 5) to stimulate the ABA response and stress tolerance in Arabidopsis (Chen et al., 2021), while the overexpression of *OsRab11* in rice (*Oryza sativa*) resulted in a decreased plant sensitivity to ABA and enhanced plant tolerance to salt and osmotic stresses (Chen, Heo, 2018). Here, we found that *FAMOSS* OE plants were less sensitive to ABA treatment and strongly inhibited by salt stress. Moreover, the stress-related proteins were downregulated in both the *FAMOSS* OE and KO lines, suggesting a link between the functions of the Rab proteins and *FAMOSS* expression levels.

In conclusion, we described the functions of a new plant SEP encoded by a lncRNA in the moss *P. patens*. We suggest that this peptide, named FAMOSS, is a possible interaction partner of the Rab GTPases, which in turn control numerous processes, including vesicular trafficking, polar tip growth, phytohormone signaling, and stress tolerance, processes that were altered in the *FAMOSS* OE/KO mutants. The *FAMOSS* peptide could therefore be considered an important component of Rab signaling in plants. Further studies will help elucidate the mechanisms of such interactions.

## Acknowledgments

This research was supported by the Russian Science Foundation (project No. 17-14-01189). We thank Igor Mazheika for the help with plant cultivation and good advices.

## Author contributions

A.M. and I.F. conceived and designed experiments. A.M. and I.F. wrote the manuscript with input from all authors. I.F. supervised the project. A.K and A.G. performed nanopore sequencing and analyzed the sequencing data. A.M., S.K. and R.Z. performed the proteomics analysis. A.M., N.G., I.S., V.R., A.N. and A.F. conducted analysis of phenotypes and fluorescent microscopy. N.G. and A.K. analyzed transcription using RT-PCR. I.F. performed the statistical and bioinformatics analyses. A.K., D.K., V.M. and V.L. created genetic construction, generate recombinant proteins, and performed moss transformation experiments. I.S., M.P. and A.M. performed pull-down and co-immunoprecipitation. All authors read and approved the final manuscript.

## Declaration of Interests

The authors declare no competing interests.

## Materials and Methods

### Physcomitrium patens growth conditions

*Physcomitrium patens* subsp. patens (“Gransden 2004”, Frieburg) protonemata of wild type and FAMOSS (PSEP1) - knockouting and overexpressing lines (Fesenko *et al*., 2019) were grown on BCD medium supplemented with 5 mM ammonium tartrate (BCDAT) and/or 0.5% glucose during a 16-h photoperiod at 25 C in 9-cm Petri dishes (Nishiyama et al. 2000). Growth rate measurements were taken at 30 day. The images of protonemal tissues and cells were obtained by a Microscope Digital Eyepiece DCM-510 attached to a Stemi 305 stereomicroscope (Zeiss, Germany) or Olympus CKX41 (Olympus, Japan) at 10 or 30 day. Seven-day-old protonema tissues grown in liquid BCDAT medium were used for proteomic, pull-down assay, co-immunoprecipitation and qRT-PCR analyses. The gametophores were grown on free-ammonium tartrate BCD medium under the same conditions, and 8-week-old gametophores were used for the further analysis.

### Quantitative reverse transcription real-time PCR

Total RNA from protonema and gametophores was isolated as previously described (Cove et al., 2009). RNA quality and quantity were evaluated via electrophoresis in an agarose gel with ethidium bromide staining. The concentration of total RNA was measured using a Quant-iT™ RNA Assay Kit (Thermo Fisher Scientific), 5–100 ng on a Qubit 3.0 (Invitrogen, US) fluorometer. The cDNA for qRT-PCR was synthesized using an MMLV RT Kit (Evrogen, Russia) according to the manufacturer’s recommendations employing oligo(dT)17 -primers from 2 μg total RNA after DNase treatment. The primers were designed using Primer-BLAST (Ye et al., 2012) (Supplemental Table 4).

### Confirmation of *FAMOSS* transcription using nanopore RNA

Nanopore data were downloaded from the BioProject with the accession number PRJNA681088 (Fesenko et al., 2021). Reads were mapped against the genome Physcomitrium patens V3.3 (Lang *et al*., 2018) by minimap 2.17 (Li, 2016) with the following parameters: -ax splice -uf -k14 f -G2k. ONT reads with a primary alignment to the genome were retained. The obtained ‘SAM’ files were sorted and indexed with ‘SAMtools’(Li et al., 2009). Sashimi plot was constructed in IGV v2.6.3 (Thorvaldsdottir et al., 2013).

### Generation of the overexpressing lines with FAMOSS-FLAG peptides

The FAMOSS sequences fused with FLAG (DYKDDDDK) tag were obtained by PCR with genomic DNA as a template and primers listed in the Supplemental Table 4. Amplicons were cloned into the pPLV27 vector (GenBank JF909480) using the ligation-independent cloning (LIC) procedure (Aslanidis and de Jong, 1990; De Rybel et al., 2011). The resulting plasmids - pPLV27-pep4-g4s-C-flag and pPLV27-N-flag-g4s-pep4 contain the C-terminus or the N-terminus FLAG-fused FAMOSS peptides respectively (Supplemental Figure 5). Before transformation, the plasmids were purified by Qiagen Plasmid Maxi Kit (Qiagen, Germany). Moss protoplasts were transformed with circular pPLV27-pep4-g4s-C-flag and pPLV27-N-flag-g4s-pep4 plasmids. Mutant lines with the overexpression of fused FAMOSS-FLAG peptides were generated as described previously (Fesenko *et al*., 2019)

### CRISPR/Cas9-directed fusion with β-D-glucuronidase (GUS) protein

The β-glucuronidase (GUS) reporter gene system was used to study spatial expression of FAMOSS peptide. The peptide coding sequence was used to search for CRISPR RNA (crRNA) preceded by a *S. pyogenes* Cas9 PAM motif (NGG) using the web tool CRISPR DESIGN (http://crispr.mit.edu/). The crRNA closest to the stop codon of FAMOSS peptide was selected for cloning. The mutant lines with FAMOSS coding sequence fused with GUS reporter protein were created using the CRISPR/Cas9 system and the pTZ-donor4 plasmid (Supplemental Methods). Protoplasts were transformed with circular sgRNA, pTZ-donor4 and pBRF as previously described (Fesenko *et al*., 2019).

### GUS activity assay

Histochemical GUS staining was carried out using β-Glucuronidase Reporter Gene Staining Kit (Sigma-Aldrich) according to the manufacturer’s recommendations. Moss tissues were vacuum-infiltrated with Staining Solution for 2 minutes and incubated at 37°C overnight (> 16 hours). Stained samples were imaged by Microscope Digital Eyepiece DCM-510 attached to a Stemi 305 stereomicroscope (Zeiss, Germany) or Olympus CKX41 (Olympus, Japan) in the bright-field.

### The conservation analysis

To identify the FAMOSS orthologs in different plant taxa, TBLASTN search with default parameters was performed using CNGBdb BLAST Service for 1000 Plants (oneKP or 1KP) trancriptomes (https://db.cngb.org/onekp/). The alignments were filtered by *E*-value <10^−3^ cut-off and manually curated.

The full peptide sequences from TBLASTN search were then aligned with MAFFT v. 7 (Katoh and Standley, 2013) and transformed into an HMM profile by hmmbuild tool from HMMER v3.3.2 package (Eddy, 2011). Orfipy tool (Singh and Wurtele, 2021) was used to predicted proteins (> 30 aa) from the transcriptomes of conifers and eudicots species that were downloaded from https://datacommons.cyverse.org/browse/iplant/home/shared/commons_repo/curated/oneKP_cap stone_2019. Hmmsearch tool (hammer.org, *E* < 0.01) was used to identify possible FAMOSS orthologs in this protein dataset and obtained hits were manually curated.

### Sequence Alignment and Conserved Motif Identification

The full amino acid sequences of the identified FAMOSS orthologs were extracted from the corresponding transcripts and used for multiple alignments using MAFFT v.7 with default settings. The resulting multiple sequence alignment (MSA) was inspected and adjusted manually. The online programs MEME (Multiple Expectation Maximization for Motif Elicitation; https://meme-suite.org/meme) with default parameter settings were used to search for conserved motifs.

### The phylogenetic analysis of the FAMOSS orthologs

We reconstructed a maximum-likelihood (ML) phylogenetic tree based on the entire MSA of 250 orthologs using IQ-TREE v.1.6.12 (Nguyen et al., 2015) with autodetected models (JTTDCMut+G4 model). Extremely divergent sequences were excluded from the analysis. The phylogenetic trees were generated by the Toytree package (Eaton et al., 2020).

### Prediction of FAMOSS 3D-structure

In order to predict FAMOSS peptide 3-D structure we used AlphaFold2 (Jumper *et al*., 2021) algorithm, which was run as Google Colaboratory project on GitHub with custom multiple sequence alignment (MSA) option. For MSA construction for a particular peptide we firstly obtained a list of 100 mostly similar sequences using BLASTP search against a file containing possible FAMOSS homologs. We then performed multiple protein alignment with M-COFFEE online tool with default parameters (http://tcoffee.crg.cat/apps/tcoffee/do:mcoffee (Notredame et al., 2000). Resulting MSA .fas file was used in the algorithm with default parameters. Structures were visualised with PyMol v.2.3.0 (https://github.com/schrodinger/pymol-open-source).

### LC-MS/MS analysis and protein identifications

Quantitative proteomic analysis was conducted as described previously (Spechenkova et al., 2021). Additional details are described in the Supplemental Methods. Briefly, proteins were extracted by phenol extraction method (Faurobert et al., 2007), digested by 1 µg sequence-grade modified trypsin (Promega, Madison, WI, USA) at 37 °C overnight, iTRAQ labelling (Applied Biosystems, Foster City, CA, USA) was conducted according to the manufacturer’s manual. Proteins were labelled with the iTRAQ tags as follows: Wild type – 113, 114, 115 isobaric tags, FAMOSS OE and KO – 116, 117, 118 ones. The samples after pull-down assay and co-immunoprecipitation were also digested by 1 µg sequence-grade modified trypsin (Promega, Madison, WI, USA) at 37 °C overnight in the same manner, but were not labeled by iTRAQ labels. LC-MS/MS analysis was carried out on an Ultimate 3000 RSLCnano HPLC system connected to a QExactive Plus mass spectrometer (Thermo Fisher Scientific, USA). Tandem mass spectra were analysed by PEAKS Studio version 8.0 software (Bioinformatics Solutions Inc., Waterloo, Canada). The custom database was built from Phytozome database *P. patens* combined with chloroplast and mitochondrial proteins (33,053 records).

### Pull-down assay

The plasmids pET-mSAV and pET-mSAV-pep4 were generated based on pET-plasmid (Novagen, USA) and contained sequence coding for mature streptavidinstreptavidine (SAV) fused with the sequences for TEV-proteinase site (pET-mSAV) or the same sequences fused with and FAMOSS peptide (pET-mSAV-pep4). The *E. coli* Rosetta 2 (DE3) strain was used to obtain recombinant proteins. Expression was induced by adding IPTG to a final concentration of 0,5 mM. The cells were disrupted by sonication using a Branson Sonifier 250 (VWR Scientific, USA) sonicator according to the manufacturer’s instructions. The lysate was purified by centrifugation (15,000× g, 25 min) and washed twice with 1% (v/v) Triton X-100. The pellet was dissolved in 6M guanidine chloride, pH 1,5, dialysed against 200 mM NaHCO3 pH 8,0 and purified using iminobiotin agarose (Thermo Fisher Scientific, USA) according to manufacturer’s instructions.

To the tissue of wild type protonema, grounded in liquid nitrogen, the lysis buffer (PBS, 150 mM NaCl, 0,1% Triton X-100, 0,2% NP-40, Protease inhibitor cocktail for plant cell extracts (Sigma)) was added. Lysate was incubated on ice for 30 min with rotation (Heidolph Duomax 1030) and centrifuged 14 000g, 15 min, 4 C (Eppendorf Centrifuge 5418 R). 15 µg of recombinant FAMOSS-streptavidin/streptavidin was added to supernatant and incubated 1h on ice with rotation. Proteins in solution were cross-linked by 3 mM DSP (Thermo Fisher Scientific Pierce) and incubated during 2 h. Reaction was quenched by 40 mM Tris-HCl рH 8 during 15 min on ice, then 50 mM ammonium bicarbonate was added. Samples were centrifuged at 14 000 g 10 min. The 20 mkl of pre-washed by washing buffer (50 mM ammonium bicarbonate-NaOH рН 11, 500 mM NaCl, 0,1% Triton X-100, 0,2% NP-40) iminobiotin agarose (Thermo Fisher Scientific Pierce) was added to supernatant and incubated on ice during 1h, washed 4 times by washing buffer. Proteins of interest were eluted three times by 50 mM ammonium acetate pH3 with 500 mM NaCl. Eluates was neutralised by 20 mM Tris and clean-uped by Amicon Ultra-0.5 Centrifugal Filter Unit Ultracel-3K (Millipore). After that the crosslinker was cleaved with 20 mM DTT by heating at 50 C.

### Co-immunoprecipitation

To identify the proteins which interact with FAMOSS peptide the co-immunoprecipitation approach was used. The plant tissue, which was grounded at liquid nitrogen, was homogenized in 450 mkl lysis buffer (50 mM Tris-HCl pH 8, 150 mM NaCl, 0,1% Tween-20, 10 mkl/ml protease inhibitor cocktail (Sigma)). The resulting suspension was incubated on ice 30 min with rotation and centrifuged at 13 000g for 15 min, 4 C (Eppendorf Centrifuge 5418 R). Proteins in the clear supernatant (protein extract) were cross-linked with 5 mM DSP (Thermo Fisher Scientific Pierce) for 2 h with agitation. The reaction was stopped with 40 mM Tris-HCl рH 8 during 15 min incubation. Samples were centrifuged at 14 000 g 10 min. The 50 µl pre-washed with washing buffer (50 mM Tris-HCl pH 8, 150 mM NaCl, 0,1% Tween-20) M2 anti-FLAG magnetic beads (Sigma) was added to supernatant. The samples were incubated 2 h on ice with rotation of the washing buffer and once with 700 mkl of TBS (50 mM Tris-HCl pH 8, 150 mM NaCl). The protein complexes were specifically eluted with 125 mkl of 100 ng/mkl 3xFLAG peptide (Sigma) solution in TBS during 30 min on-ice incubation with agitation. The crosslinker was cleaved with 20 mM DTT by heating at 50 C.

### SDS-PAGE and in-gel trypsin digestion

Proteins after pull-down and Co-IP experiments were vacuum dried, dissolved in a sodium dodecyl sulphate-polyacrylamide gel electrophoresis (SDS-PAGE) loading buffer (Bio-Rad) and separated on 15 % SDS-PAGE gels following the Laemmli method (Laemmli, 1970). Gels were stained as described previously (Schagger, 2006). The gel pieces corresponding 25 kDa MW were cut, destained by 15 mM sodium thiosulfate and 50 mM potassium ferricyanide. Protein in-gel trypsin digestion was conducted as described by (Rubtsova *et al*., 2018).

### Regeneration of protonema from protoplasts

Protoplast was prepared from protonemata as described previously (Fesenko et al., 2015) and incubated during the 6 days on solid BCD agar medium. Regenerated protonema was detected by microscopy analysis.

### Fluorescent microscopy

To detect vesicles, protonemal cells were stained by the 5 µM of fluorescent dye SynaptoGreen C4 (equivalent to FM1-43, Biotium, USA. SynaptoGreen C4 is a synonym of fluorescent probe which is originally called FM1-43 dye and is available from Biotium under the trademark name of SynaptoGreen™) during 5 min and washed. To visualize cell walls the protonema cells were stained by 10 µg/mL of propidium iodide (fluorescein diacetate, Sigma-Aldrich, USA) during 5 min. Fluorescence was detected by Axio Imager M2 microscope (Zeiss) with an AxioCam 506 mono digital camera (Zeiss) and Zen 2.6 pro software (Zeiss). Filter unit 44 FITC (λex BP 475 nm/40 nm; λem BP 530 nm/50 nm) was used for SynaptoGreen C4 fluorescence detection, Filter unit 20 Rhodamin (λex BP 546/12 nm; λem BP 575–640 nm) was used for propidium iodide.

### Statistical analysis

Statistical analysis and visualisation were made in Python v. 3.7.5(Van Rossum, Drake, 1995) using modules scipy 1.5.2 (Virtanen et al., 2020), seaborn 0.11.1 (Waskom, 2021), numpy 1.20.1, pandas 1.2. 3 (McKinney, 2012).

### Data availability

The mass spectrometry proteomic data have been deposited to the ProteomeXchange Consortium via the PRIDE (Perez-Riverol et al., 2019) partner repository with the dataset identifiers PXD028296 and PXD028342.

## Notes

### Competing Interest Statement

The authors have declared no competing interest.

